# Addition of Soil Protists Enhances Performance of Agrochemical Seed Treatments

**DOI:** 10.1101/2024.06.14.599089

**Authors:** Christopher J. Hawxhurst, Travis McClure, Daniel Kirk, Mikhael Shor, Daniel J. Gage, Leslie M. Shor

## Abstract

Chemicals are an integral part of modern agriculture, and are applied through a variety of methods. Some agrochemicals applied for crop protection function by absorption through the root before translocation to the rest of the plant. To be absorbed by the root, the agrochemical must first be transported through the soil, often by water. Some agrochemicals suffer from poor water-based soil transmission due to their chemical properties, limiting their application as a traditional seed treatment. Two such agrochemicals are chlorantraniliprole and spinosad. Soil protists are an important component of the soil microbial community. Certain soil protists have been previously shown to facilitate transport and targeted delivery of suspended particles and cells through soil and microfluidic devices. We provide practical evidence that a soil protist, *Colpoda sp*., when co-inoculated with an agrochemical seed treatment, can substantially and robustly reduce subsequent pest feeding damage compared with the agrochemical alone. Using maize (*Zea mays* L.) and fall armyworm, *Spodoptera frugiperda* (J. E. Smith, 1797) (Lepidoptera: Noctuidae), in a plant damage assay, we directly compare pest feeding damage and mortality in plants that received no additional treatment, only protists, only agrochemical, and co-inoculation of agrochemical with protists. We discover for both agrochemicals tested, the co-inoculation of protists with the agrochemical increases protection in leaves when the efficacy of the agrochemical alone declines. Protist amendment is a simple, natural, inexpensive, chemical-free, soil-based transport enhancer that thus may be widely useful in a variety of contexts including more sustainable and cost-effective integrated pest management.

**Importance:** Pest resistance, regulatory pressure, and environmental concerns are limiting many classes of pesticides which can be effectively used to protect valuable crops from pests. Other classes of pesticides, however, are limited by physical characteristics – water solubility or octane-water partition coefficient (K_OW_) fall outside the limit for an effective seed coating, or the per-unit cost is high enough to discourage broad application. Here, we provide data which supports the co-inoculation of a high value, low solubility, high K_OW_ pesticides with a naturally-occurring soil protist as a seed treatment can enhance crop protection relative to the pesticide alone. This co-formulation reduced feeding damage by up to 30% compared with the pesticide alone. Co-inoculation of crop-protecting agrochemicals with natural soil protists may be employed as a more sustainable agriculture biotechnology, enabling the use of classes of agrochemicals which may not otherwise show sufficient performance for use as a seed treatment.

## 1 Introduction

Chemicals including herbicides, fungicides, insecticides, and nematicides are commonly used in agriculture. While herbicides are commonly applied as a foliar spray, many insecticides, fungicides, and nematicides require soil application and act at, or are absorbed by, the roots following transport through the soil via water (1–3). Some agrochemicals suffer from poor soil transmission, an inability to target delivery to vulnerable, actively growing roots, and health, safety, and environmental concerns from high application rates. Further, due to their high specific cost, bulk application of agrochemicals can be cost prohibitive.

Conventional application methods include broadcasting of granular agrochemicals (with pre-plant tilling or post-plant), foliar sprays, seed applications, and root drenches of liquid suspensions. Performance of granular formulations are enhanced by mixing through the upper 10 cm of soil, but this is not possible for mid-season application, treatment of annuals, no-till, intercropping, or agroforestry systems. Agrochemicals spread from granules but may not reach the root tips, especially as the plant grows. Root drenches promote delivery along the full length of root systems, but the high application rates required come with higher costs and the risk for adverse environmental impacts. Without targeting capability, the entire soil volume must be treated despite the relatively small proportion of total soil volume that is comprised by the rhizosphere. Foliar sprays enable the direct treatment of at-risk leaves, but can fail to provide translocation, which is necessary to protect new leaves (4). Applications coated over seeds prior to planting are referred to as seed treatments. This placement of agrochemicals onto the seed enables their close proximity to emerging roots, but growing roots may quickly extend beyond the zone of protection (5–7).

Water-mediated transport of agrochemicals through soil has several disadvantages including poor transmission, especially for hydrophobic agrochemicals, and the potential for wasted chemicals and adverse environmental impacts caused by runoff with excess rainfall. Mathematical modeling studies of chemical fate and transport in soils generally focus on passive, water-mediated mechanisms (2, 3, 8), despite the fact that other mechanisms exist, especially in the absence of percolating water (9). Some agrochemicals show low crop absorption for seed treatments, with overall recovery below 1.5% of the applied agrochemical (10). Facilitating transport and targeting delivery of agrochemicals in the rhizosphere could improve agrochemical efficacy at lower application rates, reducing both overall treatment cost and adverse environmental impacts by runoff by keeping the agrochemical near the crop roots for longer (11–15). Facilitating transport and targeting delivery may also enable the use of additional classes of agrochemical as seed treatments. This allows for additional “tools” for use against pests, which helps avoid regulatory limits and helps prevent the development of pest resistance in the field (16).

We have previously shown soil protists facilitate transport and target delivery of suspended particles (17, 18). Our prior work focused on direct observations of soil-based processes and building mechanistic understanding. In Rubinstein et al., we quantified the extent of protist-facilitated transport in emulated soil micromodels and described three distinct mechanisms of protist-facilitated transport: translocation of (*i*) ingested and (*ii*) surface-attached fluorescent particles, and (*iii*) movement of suspended particles in protist-engendered fluid flow streams (17). In a subsequent report, we showed direct photomicrographic evidence for enhanced spatial distribution of live fluorescent bacteria, despite feeding pressure, by the addition of protists. We further demonstrated high bacterial abundance along roots, including root tips, especially in low moisture conditions (18). Here we show treating seeds with formulated insecticides plus protists substantially and robustly reduces subsequent feeding damage in a maize leaf fall armyworm damage assay compared with similar plants treated with insecticides alone. This report thus provides practical evidence for systemic benefits of co-application of soil protists. Seed treatments that also contain soil protists thus may enable lower therapeutic dosages in a variety of application methods (i.e., seed treatments, surface-applied granules, or low volume soil applications) versus similar insecticide products formulated without protists. Protist amendment is a simple, natural, inexpensive, chemical-free, soil-based transport enhancer that thus may be widely useful in a variety of contexts including more sustainable and cost-effective integrated pest management.

## 2 Results

### 2.1 Overall Feeding Damage

Overall feeding damage was computed by averaging, for each plant, the damage measures for each of the six leaf segments. Results by treatment are plotted in Figure 1 with different colors within each panel reflecting significant differences in means (p < 0.001). No significant difference in mean overall feeding damage was found between Baseline and Protists treatments (94% vs. 95% feeding damage, p > 0.10). Both the Chlorantraniliprole treatment and the Spinosad treatment exhibited significantly less mean overall feeding damage (62% and 53%, respectively) than either Baseline or Protists treatments (p < 0.001). Co-application of protists along with each agrochemical further reduced feeding damage (p < 0.001). Mean overall feeding damage was 53% in Chlorantraniliprole + Protists and 32% in Spinosad + Protists.

**Figure 1.**
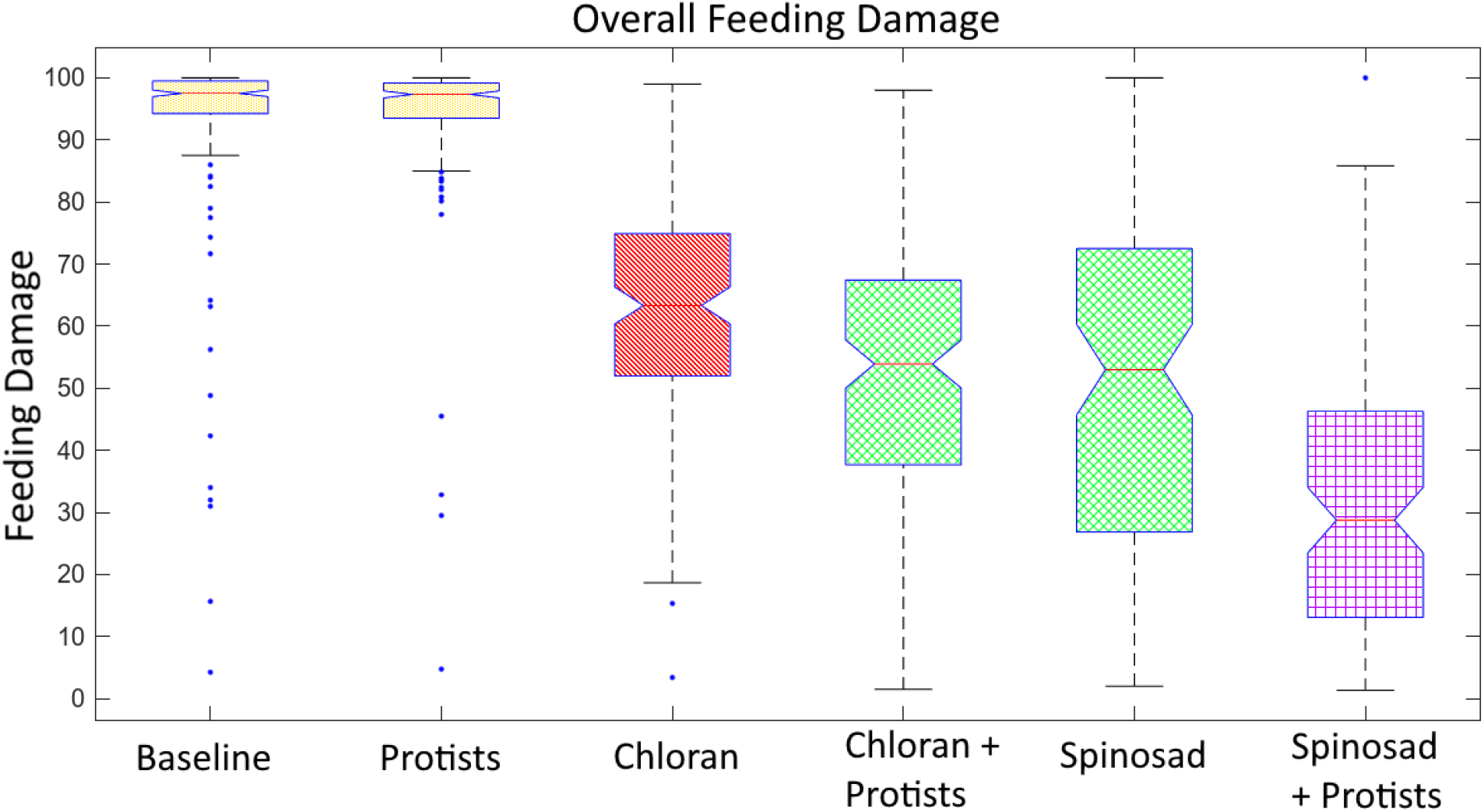
Box-and-whisker plot of feeding damage by treatment across all data points. Boxes contain the middle 50% of data points, and the median is marked by a red line with a notch. Individual plots are grouped by color using Bonferroni-corrected pairwise t-tests – treatments colored similarly imply p > 0.05 (although all are p > 0.10). Different colors imply p < 0.05 (although all are p < 0.001). Treatments containing agrochemicals (Chlorantraniliprole, Chlorantraniliprole + Protists, Spinosad, Spinosad + Protists) are statistically distinct compared with treatments without agrochemicals (p < 0.001). Treatments without agrochemicals (Baseline, Protists) show no significant differences (p > 0.10).

Data for Baseline and Protists were pooled across all 5 experimental runs while the remaining are only available within subsets of the runs (experimental runs 1, 2, 3 for Chlorantraniliprole; runs 4, 5 for Spinosad). Statically similar results were obtained when restricting Baseline and Protists data pools to similar subsets (Runs 1, 2, 3 or Runs 4, 5). The regressions confirmed the above results: no significant difference between Baseline and Protists (p = 0.433), and highly significant differences (all p < 0.001) between Baseline and either agrochemical, Protists and either agrochemical, and between each agrochemical with and without Protists. Further robustness tests involved using average FAW mortality across the six measurements for each plant instead of damage as the independent variable (Table SI 2). Again, no significant difference between Baseline and Protists was observed (p = 0.472) and highly significant differences (all p < 0.001) for other treatment comparisons.

In summation, Chlorantraniliprole + Protists led to 44% less feeding damage than Baseline or Protists and 15% less feeding damage than Chlorantraniliprole alone (62% to 53%). Spinosad + Protists led to 66% less feeding damage than Baseline or Protists and 40% less feeding damage than Spinosad alone (53% to 32%). These reductions are all highly statistically significant (p < 0.001).

### 2.2 Chlorantraniliprole by Leaf Segment

Treatment with Chlorantraniliprole was analyzed based on how different leaf segments were protected from herbivory. Feeding damage for each leaf segment for Chlorantraniliprole with and without protists are shown in Figure 2, with Baseline and Protists treatments included for comparison. In all cases, the Chlorantraniliprole + Protists treatment had significantly less feeding damage than either Baseline or Protist treatments (p < 0.001). The effect of Chlorantraniliprole without protists varied by leaf number with early leaves being better protected than later leaves. Chlorantraniliprole significantly reduced feeding damage relative to Baseline and Protist treatments (p < 0.001) in both segments of Leaf 3 and in Leaf 4 Tip and Leaf 5 Tip with the remaining two segments (Leaf 4 Middle, Leaf 5 Middle) showing no or only marginal significant reductions (p ≥ 0.050).

**Figure 2.**
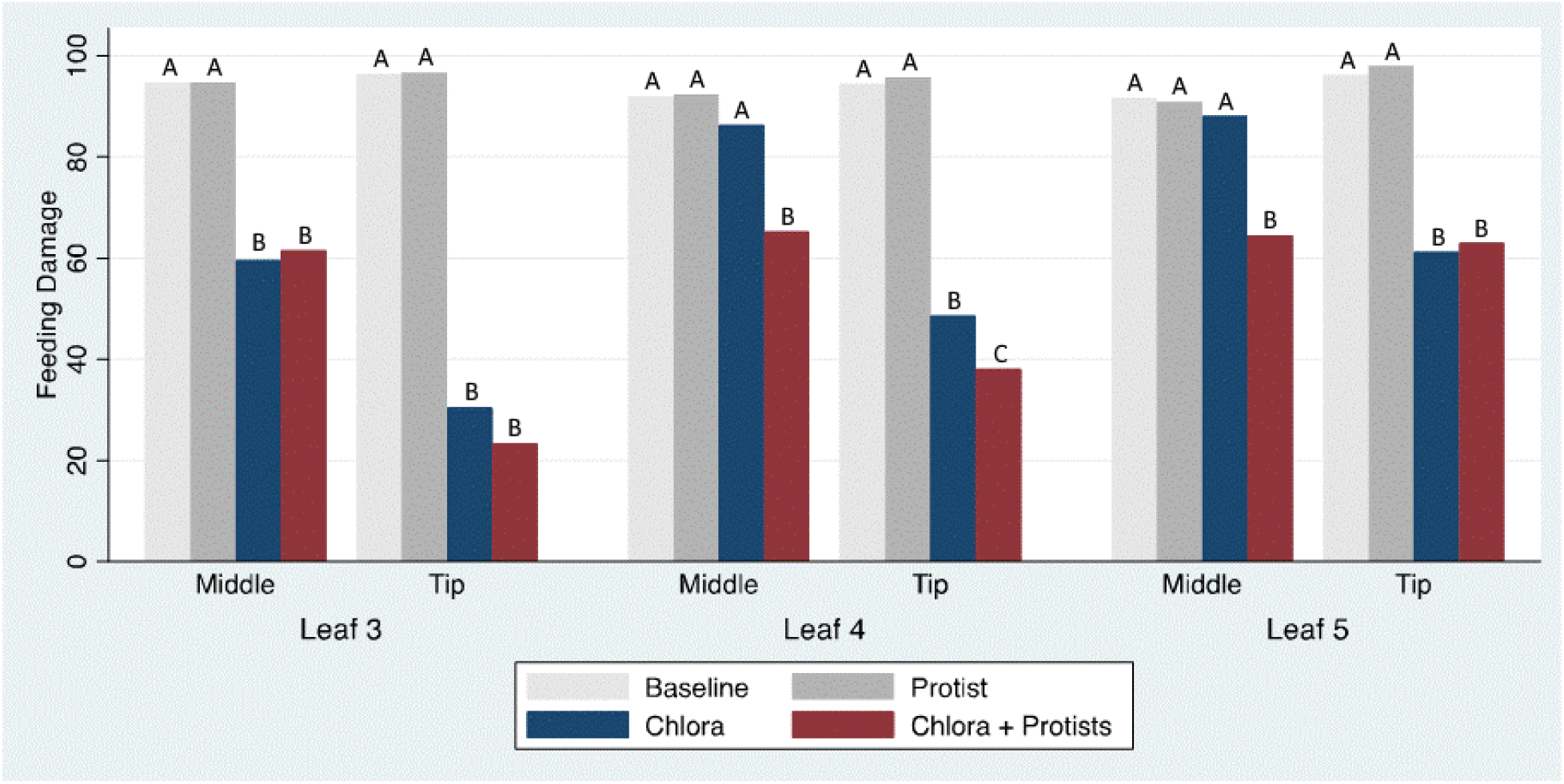
Feeding damage following treatment with Chlorantraniliprole, separated by leaf and section. Letters indicate statistical significance, where treatments with the same letter are statistically similar (p ≥ 0.050), while treatments with different letters are statistically different (p < 0.050). Specifically, Chlorantraniliprole treatment shows a statistically significant difference from Baseline and Protist treatments in both segments of Leaf 3, Leaf 4 Tip, and Leaf 5 Tip (p < 0.001). In all cases, the Chlorantraniliprole + Protists treatment had significantly less feeding damage than either Baseline or Protist treatments (p < 0.001).

Comparing feeding damage between Chlorantraniliprole and Chlorantraniliprole + Protists treatments, the results again varied by leaf number. Chlorantraniliprole + Protists significantly reduced feeding damage relative to Chlorantraniliprole for Leaf 4 Tip (49% vs. 38%, p = 0.013), Leaf 4 Middle (86% vs. 65%, p < 0.001), and Leaf 5 Middle (88% vs. 64%, p < 0.001). The co-application of protists to Chlorantraniliprole was not additionally protective (p > 0.050) for either segment of Leaf 3 or for Leaf 5 Tip. Importantly, in both segments where Chlorantraniliprole alone did not significantly reduce feeding damage (Leaf 4 Middle and Leaf 5 Middle), the co-application of protists significantly reduced feeding damage.

Leaf 4 Middle and Leaf 5 Middle show the largest difference in feeding damage reduction between Chlorantraniliprole and Chlorantraniliprole + Protists. For Leaf 4 Middle, Chlorantraniliprole reduces damage relative to Baseline from 93.6% to 86.4%, a 7.2 percentage point reduction. Chlorantraniliprole + Protists reduces damage to 65.2%, a reduction of 28.4 percentage points from Baseline, or nearly 4 times the benefit of Chlorantraniliprole alone. Similarly, Leaf 5 Middle sees damage reduction of 3.5 percentage points from Chlorantraniliprole relative to baseline while Chlorantraniliprole + Protists reduces damage by 27.2 percentage points, or more than 7 times the benefit of Chlorantraniliprole alone.

### 2.3 Spinosad By Leaf Segment

For the Spinosad treatments, feeding damage by leaf segment displayed the same trends as overall feeding damage (all feeding damage grouped by plant), with some slight differences (Figure 3). In almost all cases, Spinosad showed highly significantly lower feeding damage than either Baseline or Protists (p < 0.001 pairwise t-tests with Bonferroni adjustment for multiple hypothesis tests) except for Leaf 5 Middle which exhibited lower significance (p = 0.014 relative to Protists, p = 0.053 relative to Baseline). Unlike Chlorantraniliprole, however, there is a statistically significant difference between Spinosad alone and Spinosad + Protists for each leaf segment (p < 0.001).

**Figure 3.**
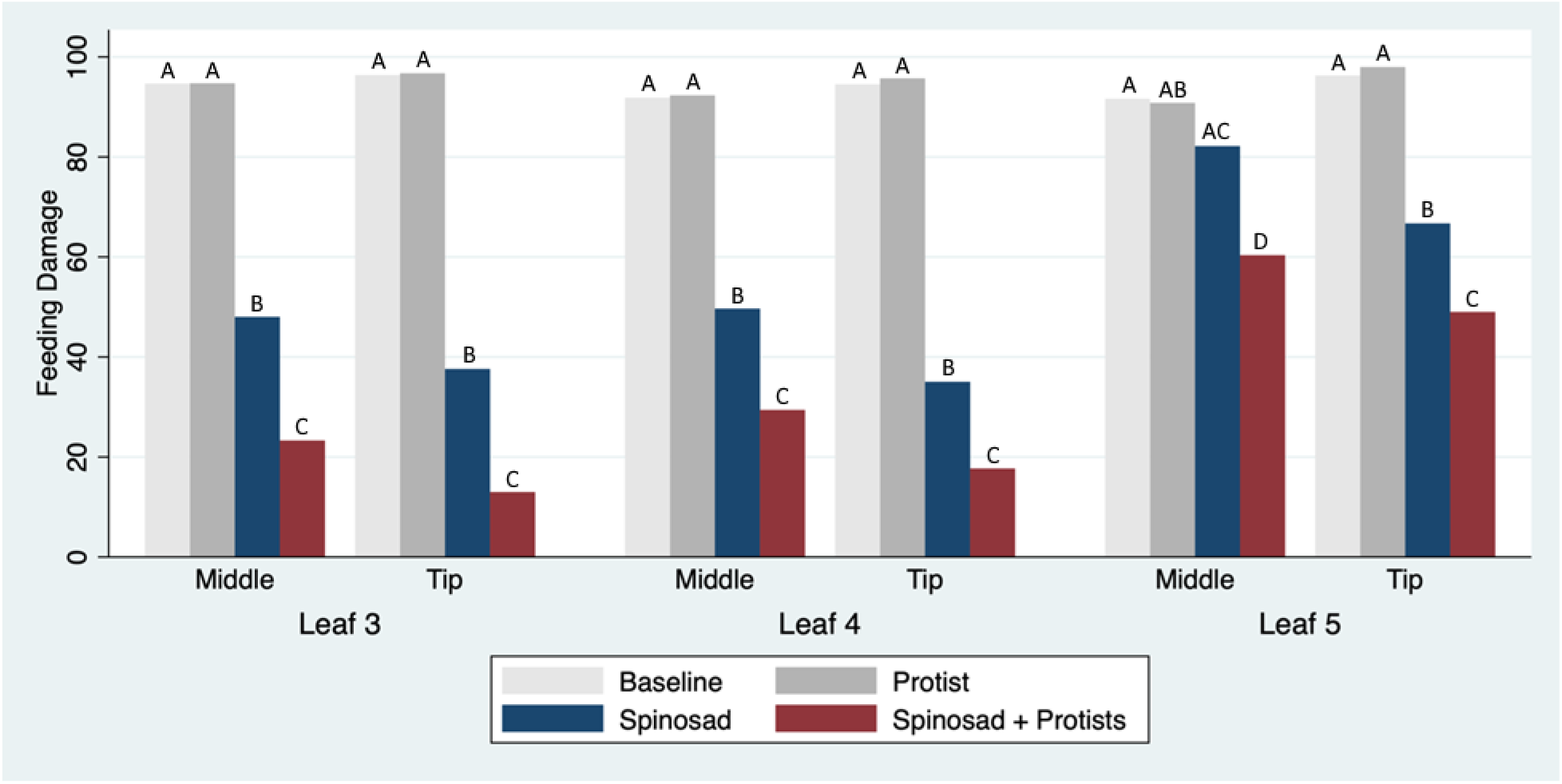
Feeding damage following treatment with Spinosad separated by leaf and section. Letters indicate statistical significance, where treatments with the same letter are statistically similar (p ≥ 0.050), while treatments with different letters are statistically different (p < 0.050). In the case of Leaf 5 Middle, the Spinosad treatment (AC) is statistically different from the Protist treatment (AC, p = 0.014), but does not meet the threshold for statistical significance from Baseline (A, p = 0.053). In all other cases, Spinosad treatment shows a statistically significant difference from Baseline and Protist treatments. Feeding damage for Spinosad treatment is statistically different from Spinosad + Protists in all leaf segments.

Because Spinosad was more effective than Chlorantraniliprole in reducing damage even without co-application of protists, the relative effect of adding protists is less dramatic with Spinosad than with Chlorantraniliprole. Leaf 4 Tip saw the lowest feeding damage from Spinosad alone, reducing feeding damage by 58.5 percentage points relative to Baseline (93.5% vs. 35.0%). Spinosad + Protists resulted in a 75.8 percentage point reduction (to 17.7%), or a 30% improvement in feeding reduction over Spinosad alone. For Leaf 5 Middle, where Spinosad alone was least effective, Spinosad reduced feeding damage relative to Baseline by 9.4 percentage points (91.6% vs. 82.2%) while Spinosad + Protists resulted in more than three times the reduction, 31.3 percentage points (to 60.3%).

### 2.4 Protist Benefit

To isolate the effects of co-application of soil protists with each agrochemical, each agrochemical treatment was compared with its respective agrochemical + protists treatment in a regression framework controlling for leaf and leaf section (Table 1). In columns (1) and (2), the dependent variable is log of feeding damage, so that coefficients can be interpreted as the percentage change in damage.

**Table 1.** Regressions of logged feeding damage and mortality on leaf sections and the co-application of protists. Columns (1) and (3) use data from the Chlorantraniliprole and Chlorantraniliprole + Protists treatments. Columns (2) and (4) use data from the Spinosad and Spinosad + Protists treatments.

|  | (1)<br>Log(damage) OLS<br>Chlorantraniliprole | (2)<br>Log(damage) OLS<br>Spinosad | (3)<br>Mortality (logistic)<br>Chlorantraniliprole | (4)<br>Mortality (logistic)<br>Spinosad |
| --- | --- | --- | --- | --- |
| Protist co-application | -0.303***<br>(0.073) | -0.772***<br>(0.131) | 0.594***<br>(0.138) | 0.984***<br>(0.227) |
| Middle Section | 0.847***<br>(0.042) | 0.530***<br>(0.045) | -1.925***<br>(0.123) | -0.852***<br>(0.094) |
| Leaf 4 | 0.414***<br>(0.055) | 0.432***<br>(0.079) | -0.845***<br>(0.123) | -0.635***<br>(0.115) |
| Leaf 5 | 0.751***<br>(0.059) | 1.236***<br>(0.081) | -0.968***<br>(0.164) | -1.798***<br>(0.154) |
| Constant | 2.961***<br>(0.069) | 2.636***<br>(0.120) | 0.073<br>(0.132) | 0.545**<br>(0.192) |
| Observations | 1716 | 1150 | 1716 | 1150 |
| R<sup>2</sup> / Pseudo R<sup>2</sup> | 0.212 | 0.250 | 0.159 | 0.138 |

*Leaf 3 and Tip section are the omitted reference categories.*
*Robust standard errors clustered by plant are in parentheses.*
*P-values for F test and Wald chi-squared test for regression and logit, respectively, are $p < 0.001$ .*
*\*\*\* $p < 0.001$ , \*\* $p < 0.010$ , \* $p < 0.050$*

We again observe higher feeding damage in later leaves, with Leaf 5 exhibiting higher damage than Leaf 4, and Leaf 4 exhibiting higher damage than Leaf 3, the reference category. We also observe higher damage in the Middle sections of leaves than in the Tip sections. However, in aggregate, we observe significant reduction of feeding damage due to co-application of protists for each agrochemical, reducing feeding damage by 30% when co-applied with Chlorantraniliprole than when Chlorantraniliprole is applied alone, and reducing feeding damage by 77% when co-applied with Spinosad. These reductions are highly statistically significant (p < 0.001). Columns (3) and (4) of Table 1 serve as a robustness check by using FAW mortality as the dependent variable in a logistic regression. The same results are observed, with coefficients flipped in sign since FAW mortality is associated with lower feeding damage.

## 3 Discussion

Based on the feeding damage data gathered from the experiments described here, we conclude that the addition of protists to a seed treatment with Chlorantraniliprole or Spinosad can improve both the protection provided by, and the duration for which the agrochemical offers protection to the plant. This is best demonstrated by the middle sections of Leaf 5 – the last leaf to emerge. For both agrochemicals tested – Chlorantraniliprole and Spinosad – the addition of protists to the agrochemical offered increased protection against pests (reduction in feeding damage) compared with the agrochemical alone. Chlorantraniliprole offers a particularly interesting case – the Chlorantraniliprole alone does not offer any statistically significant protection by the Middle segment of Leaf 5, while co-inoculation of Chlorantraniliprole + Protists increases the amount of protection offered (decreased feeding damage) when compared with Chlorantraniliprole alone, extending the protection into leaf segments that are outside the zone of protection of Chlorantraniliprole alone. It is worth noting, however, testing was performed in pasteurized soil for consistency, and living soil may affect the results.

While we lack direct evidence of protists actively transporting agrochemicals throughout the root network, we have secondary evidence that transportation is the method by which Protists enhance the protection offered by the agrochemicals tested here. Leaf number should correlate with root network size, i.e., the root network was smaller when the earliest leaf tested (Leaf 3) was formed compared to its size when the last leaf (Leaf 5) was formed (Figure 4). Looking at the results for these leaves, the addition of protists to a seed treatment containing agrochemical did not seem to confer significant benefits for Leaf 3 which developed with the smallest average root network and with the longest time to accumulate agrochemical. However, leaves with larger average root network size, such as Leaves 4 and 5, did show statistically significant benefits from the co-inoculation of Protists with the agrochemical compared with the agrochemical alone. This secondary evidence combined with previous research on protist-based transportation (17, 18), we can conclude that Protists can transport the agrochemicals tested. Further research is required to determine the extent to which Protists can aid the distribution of agrochemicals. Additionally, future testing additional leaves (e.g., Leaf 6, 7, or 8) may show a similar break where Spinosad alone offers no protection to the leaves while co-inoculation of Spinosad + Protists continues to protect from feeding damage.

**Figure 4.**
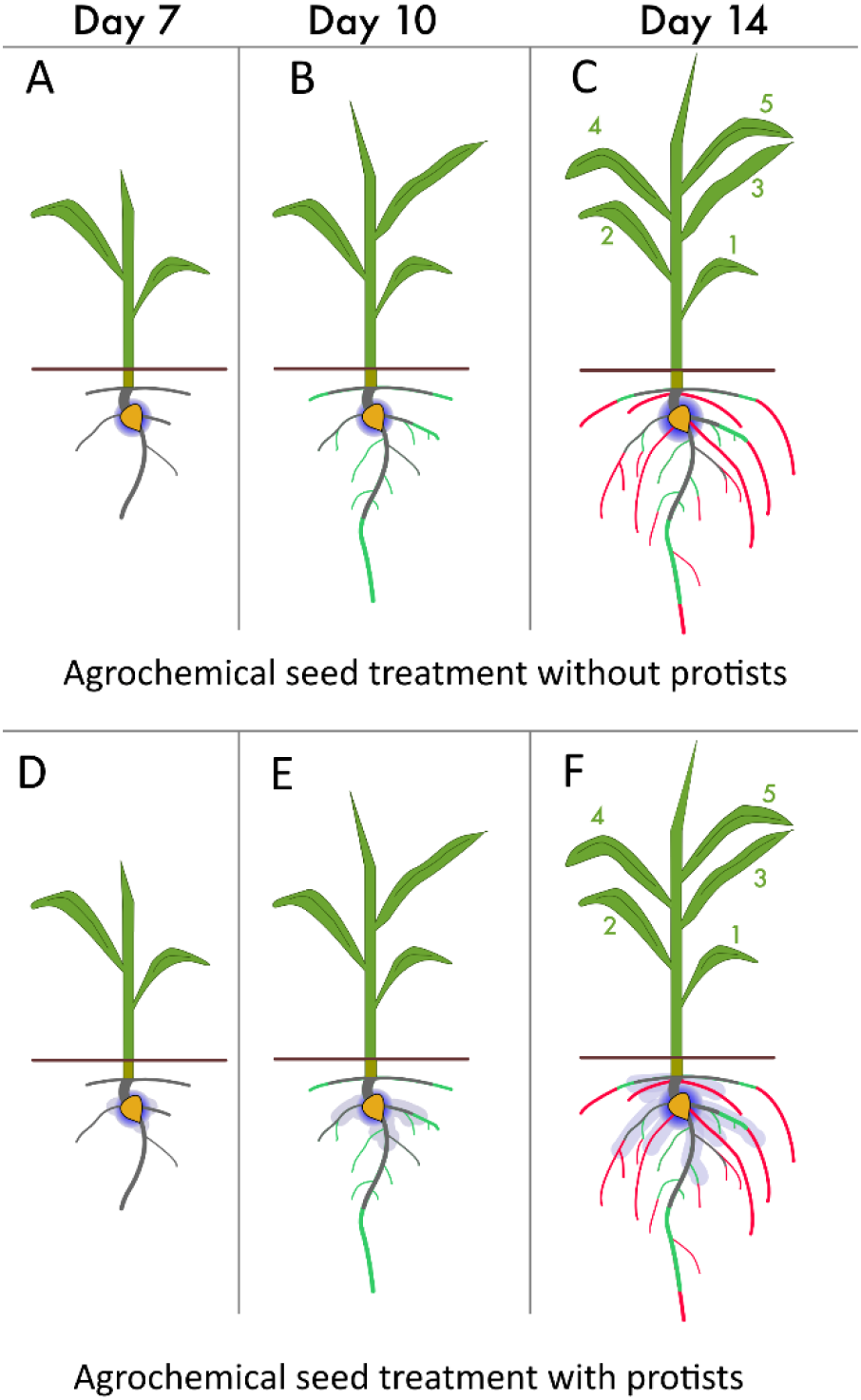
Root system expansion and delivery of insecticide to leaves following root-treatment. Seeds are treated during planting and the treatment diffuses away from seed to a small extent and provides a limited region of effective concentration (bright blue gradient and light blue extensions). Uptake of the treatment into a transpiring leaf is dependent on the surface area of the root system that is plumbed to that leaf and exposed to effective concentrations of the treatment. The figure shows approximate plant development from days 7 (A, D), 10 (B, E), and 14 (C, F). Gray roots-present at day 7 and plumbed to transpiring leaves 1 and 2. Green roots consist of new roots and extensions of pre-existing roots that grew between days 7 and 10. Some of these are plumbed to leaf 3. Red roots: this consists of new roots and extensions of pre-existing roots that grew between days 10 and 14. Much of this new tissue is outside the zone of effective treatment concentration. Some of this new tissue is plumbed to leaves 4 and 5 which matured between days 10 and 14. A, B, C show root exposure to treatment without the addition of protists. D, E, F depict a hypothesized increase in root system exposure to treatment because of transport by protists.

## 4 Conclusions

Other benefits of inoculation with soil protists may include re-establishing protists depleted by fertilization (29), regulating bacterial communities by top-down selective grazing (30), and modulating soil fertility (31). Due to the size of protists, the transport payloads would by necessity be limited to micro- and nano-particles. Agrochemicals are already milled to the micron range as part of their commercial formulations, this would not be a limiting factor. Seed applied agrochemicals are often chosen due to their high specific activity (low required therapeutic dosage), targeted treatment application, long term soil retention/treatment duration, and high cost and reduction of material over the field. When combined with a targeted transport technology, regular seed treatment could offer improved efficacy relative to seed treatment alone due to the expansion of the therapeutic zone to include more of the growing roots. The technology described here provides a potential straightforward method to boost the performance of agrochemicals, potentially enabling the use of new classes of agrochemicals as seed treatments, or reduction of the quantity required for seed treatments. This could provide economic benefits, and offer additional tools to combat the growing spread of pests to new regions.

## 5 Material and Methods

### 5.1 Experimental Overview

We designed an experiment to measure the systemic effects on pest feeding damage for plants grown from seeds treated with insecticides plus soil protists versus similar plants treated with insecticides alone. Our experiments consisted of two stages: the growth stage and the pest stage. In the growth stage, maize seeds were treated individually with one of six mixtures containing bacteria, protists, and or/agrochemicals. All treatments included heat-killed bacteria, the food source for protists. Additional potential components of each treatment were one of two agrochemicals, or soil protists, or one of two agrochemicals plus soil protists (Table 2). Treated seeds were planted and grown in soil-filled pots in a greenhouse. Plants were grown in surplus quantities and down-selected to 48 representative plants per treatment for each of 5 experimental runs (Table 3).

**Table 2.** Experimental treatments are defined by the composition of additives applied to the seed at the growth stage. Seed treatment components are expressed on a per plant (single seed) basis.

| Experimental Treatment | Constituents Added to Seed |  |  |
| --- | --- | --- | --- |
|  | Bacteria | Protists | Agrochemical |
| Baseline | $6 \times 10^7$ | - | - |
| Protists | $6 \times 10^7$ | $5 \times 10^4$ | - |
| Chlorantraniliprole | $6 \times 10^7$ | - | 0.5 μg Chlorantraniliprole |
| Chlorantraniliprole + Protists | $6 \times 10^7$ | $5 \times 10^4$ | 0.5 μg Chlorantraniliprole |
| Spinosad | $6 \times 10^7$ | - | 167 μg Spinosad |
| Spinosad + Protists | $6 \times 10^7$ | $5 \times 10^4$ | 167 μg Spinosad |

**Table 3.** Experimental design showing the number of plants harvested per treatments per experimental run. From each harvested plant, six leaf segments were challenged individually in the pest stage for a total of 5,750 segments (i.e., 960 plants × six segments per plant) tested overall.

| Experimenta<br>l Run | Treatment |  |  |  |  |  | Sample<br>d Plants |
| --- | --- | --- | --- | --- | --- | --- | --- |
|  | Baselin<br>e | Protist<br>s | Chloran-<br>traniliprol<br>e | Chloran-<br>traniliprol<br>e +<br>Protists | Spinosa<br>d | Spinosa<br>d +<br>Protists |  |
| 1 | 48 | 48 | 48 | 48 | - | - | 192 |
| 2 | 48 | 48 | 48 | 48 | - | - | 192 |
| 3 | 48 | 48 | 48 | 48 | - | - | 192 |
| 4 | 48 | 48 | - | - | 48 | 48 | 192 |
| 5 | 48 | 48 | - | - | 48 | 48 | 192 |
| Total Plants | 240 | 240 | 144 | 96 | 144 | 96 | 960 |

In the pest stage, three individual leaves were harvested from each plant, then the Tip and Middle segments of said leaves were collected. Each of the six leaf segments from the plant were placed in individual tray wells and challenged by a single insect larva, as explained further below.

### 5.2 Biological Materials and Methods

Bacteria were used to feed and propagate soil protists, as described previously (18). Briefly, *E. coli* DH5α was grown to stationary phase in Luria broth (LB) before being washed three times and resuspended in Page’s Saline. Concentration of the *E. coli* stock solution was taken using OD_600_ measured in a 100 µL suspension in a 96-well microtiter plate and were used to calculate the amount of stock solution to feed to the protists. *E. coli* stock was then heat killed at 95 °C for 1 h before use in seed treatments or to propagate protists.

Protists were identified by 18S rRNA gene sequencing as belonging to a ciliate genus, *Colpoda*, which we classify as “UC-1” (19), and were passaged from cultures originally isolated from soil (17). UC-1 cultures were maintained by removing encysted protists from culture flask surfaces (Fisherbrand™ Cell Scrapers) and passaging half an existing culture into 50 mL fresh Page’s Saline in a 175 cm^2^ culture flask (Thermo Scientific™ BioLite™ Cell Culture Treated Flasks). Protists were fed heat-killed *E. coli* DH5α which was added twice a week to a final OD_600_ = 0.025 (total feeding OD_600_ = 0.050 per week), measured as described above. Protists were concentrated and prepared for experimentation following methods for trophozoites previously described (18).

*Spodoptera frugiperda*, commonly called “Fall armyworm” (FAW) (Lepidoptera: Noctuidae, JE Smith, 1797) are among the Food and Agriculture Organization of the United Nations top 10 global pests (20). FAW larva cause crop damage by consuming foliage. As of 2016, FAW have become established in Africa where they are expected to do nearly $10B annually in crop damage (21). In the southeastern United States, Brazil and India, insecticides are often used to control FAW. But, as FAW larva feed deep within the whorl of young maize plants, foliar insecticides require high application rates to achieve adequate penetration. FAW larva were used in their second instar (growth stage) and were sourced from the insectary of Corteva Agriscience (Indianapolis, IN).

Corn (U.S.) or maize (U.K. and worldwide) (*Zea mays* L.) was chosen for demonstrating the technology because it is a fast-growing annual crop, an essential food staple, and has a huge global market. For this study a commercial hybrid was selected (Prairie Hybrid 3773). Imbibition was omitted because seeds were planted in excess and 48 uniform representative plants for each treatment were down-selected prior to leaf harvest from 60 seeds initially started per treatment per experimental run.

### 5.3 Agrochemical Selection

Chlorantraniliprole is a synthetic insecticide used against various types of insects including beetles (Order: Coleoptera), and caterpillars (Order: Lepidoptera) which includes FAW. Chlorantraniliprole was registered in 2008 in the U.S. by E.I. du Pont de Nemours and Company, Inc., for commercial use in crop protection and for treatment of landscaped areas (22). Global sales of chlorantraniliprole reached $1725 million in 2021 with an expected annual growth of 4.4% through 2027 (23). Spinosad is a natural product insecticide used against a variety of insect pests including caterpillars, flies (Order: Diptera), beetles, and locusts (Order: Orthoptera). Spinosad is a natural fermentation product by the bacteria *Saccharopolyspora spinosa*, originally isolated in 1985, composed of *spinosyn A* and *spinosyn D*. Spinosad has been approved for use in over 200 countries and the global, end-user, market size was $422 million ad of 2017 and growing (24).

We focused this study on agrochemicals that are industrially important, commonly used, and also that have features that make them likely to benefit from co-application with soil protists. Thus, we considered agrochemicals that are capable of plant phloem or xylem transport in order to exhibit systemic (i.e., whole plant, versus localized) benefits from seed treatment or soil application. Also, we considered agrochemicals with long soil half-lives > 50 d because these would remain effective during the main growth stage of commercially important annual crops without requiring re-treatment. Next, we identified agrochemicals with a certain range of physiochemical properties. When considering water-mediated transport, root-applied agrochemicals tend to be most effective when water solubility and hydrophobicity (i.e., log K_ow_) are within a preferred range (Table 4 Agrochemicals *less* soluble and/or *more* hydrophobic than the preferred range tend to exhibit poor soil transmission via water infiltration and thus are more likely to exhibit protist-facilitated transport effects. Further, we reasoned that agrochemicals available in a particulate suspension may be most readily transported by ciliates given their particle-capturing feeding behavior (17). Finally, we screened candidate formulations for operational toxicity to UC-1 cultures (i.e., live protists in a concentrated formulation should remain viable and actively moving). Chlorantraniliprole and Spinosad satisfied all criteria and were selected for study. Both agrochemicals were used in their commercial-like formulations, which is a proprietary blend containing biocides and other ingredients needed to keep the agrochemical suspended and maintain performance.

**Table 4.** Physiochemical properties for water-based transport of agrochemicals used in this study. Note Spinosad is a mix of two different spinosoids (spinosyn A and spinosyn D). Values for Spinosad are those reported by Dow-Agriscience for the mixture.

| Physiochemical Property | Preferred Range | Chlorantraniliprole | Spinosad* |
| --- | --- | --- | --- |
| Water Solubility (ppm) | 50 – 600 | 2.5 (25) | 2 (25) |
| log K<sub>ow</sub> | 0.75 – 2.5 (26) | 721 (22) | 3.2 (27) |

### 5.4 Seed Treatment Formulations

The dosage of each insecticide employed (Table 2) was determined based on previous experimental results and was intended to exhibit a “break point,” or measurable reduction in protection, within the scope of the experiment for the agrochemical (alone) treatments (i.e., to show low feeding damage in Leaf 3 and higher feeding damage in Leaf 5 for the “Chlorantraniliprole” and for the “Spinosad” treatments). These dosages are lower than the typical therapeutic dosage and were selected to best show the effects, if any, of the addition of protists to the seed treatment on subsequent pest protection.

Formulations for seed treatments (Table 2) were prepared in Page’s Saline 24 h prior to inoculation and held at 23 °C in darkness. Prior to application, each formulation was vortexed for 30 sec then a small aliquot (50 µL) was added directly on top of each exposed seed during planting. Based on our prior experience, this small-volume-aliquot performs similarly to more conventionally formulated seed coatings, enabling early-stage screening of experimental formulations that are available only in limited quantities.

### 5.5 Growth Stage

For each experimental run, 240 pots (10-cm, TLC Square Form Pots, HC Companies) were first filled with dry, pasteurized soil (Kalamazoo, Michigan 42.30 °, -85.76) CoB—Coloma loamy sand). The soil was allowed to come to equilibrium with the automated watering system from a single rail boom irrigation system (Cherry Creek Systems) in the Corteva Agriscience greenhouse 306-D1 located in Indianapolis, IN for at least 4 d prior to planting. Next, 3.8 cm deep, 2 cm wide holes were created in the center of each soil-filled pots, and a single seed was placed tip cap side down 3.8 cm below the soil surface. Each seed was immediately inoculated with a seed treatment (described above, n = 60 per treatment). Once inoculated, maize seeds were covered in soil and supplemented with 0.9 grams of phosphorus (Triple Super Phosphate 0-46-0, Easy Peasy). Treated pots were then randomized in the greenhouse to reduce systematic error (Figure 5A). The greenhouse averaged 23 °C (over the course of the trials ranged between 19 °C at night and 30 °C during the day). The plants received 100 mL water, split over 49 passes, during the first week after planting, and 200 mL water split over 98 passes during the second week after planting. During the second week, plants were also supplemented with 90 mL of fertilizer (LX 17-4-17, Jack’s Professional).

**Figure 5.**
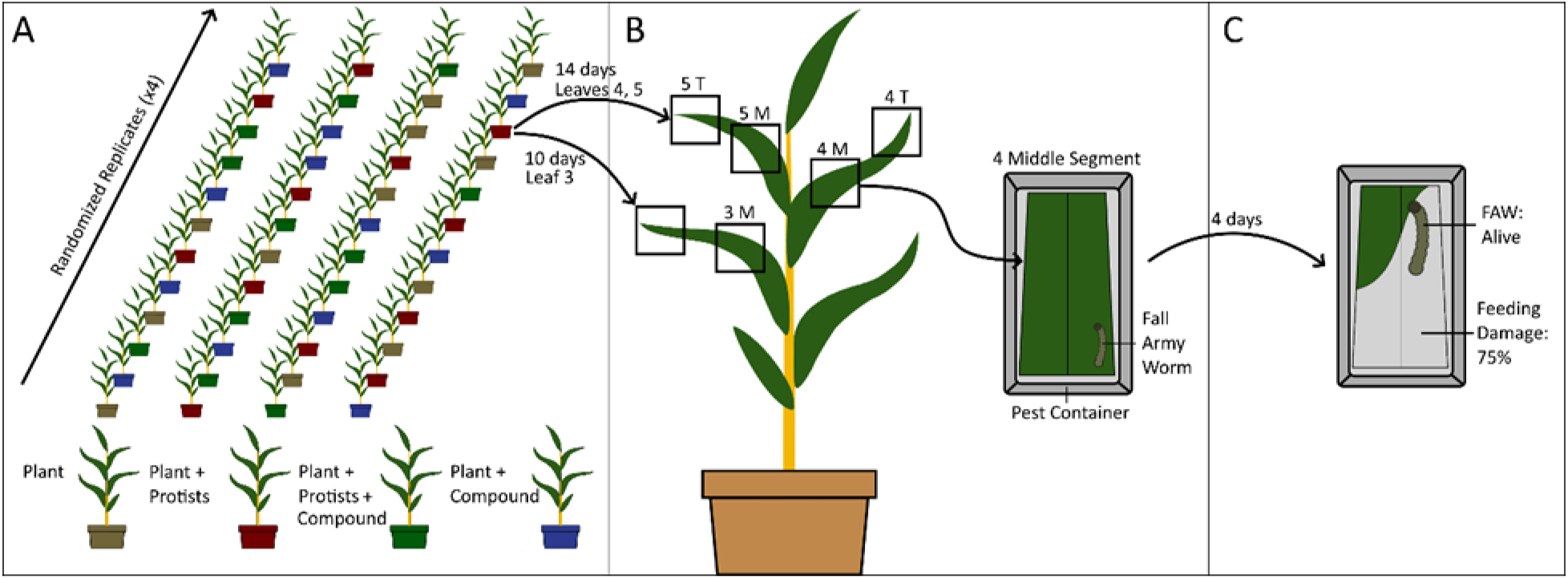
A. “Growth Stage” experimental setup showing the randomized placement of pots with 48 representative plants were selected per treatment per experiment. B. Harvest and segmentation showing (inset) the “Pest Stage”. C. Pest Stage: Evaluation assay showing ∼75% damage caused by one FAW larvae.

### 5.6 Leaf Harvest

Leaf segments were harvested on 48 representative plants selected from the 60 pots per treatment (Figure 5A). Trays were filled in a semi-random fashion with less optimal plants at the end of tray 3 and the most underdeveloped plants being discarded. Plants were numbered as they were selected during the Leaf 3 harvest, and tracked through the harvest of Leaves 4 and 5 for robustness. Maize leaves are labelled according to the order they sprout. Maize was allowed to grow for 10 d before harvesting Leaf 3 and left for another 4 d (14 d total) before harvesting Leave 4, 5 (Figure 5B). Leaves were segmented with the distal 3.8 cm labelled as Tip. Leaf Middle segments were harvested as a 3.8 cm segment from the middle of the leaf. For smaller leaves (such as the earlier-harvested Leaf 3), the Middle abutted the Tip. For larger leaves (4^th^ and 5^th^), any section between Tip and Middle segments was discarded. For agrochemicals which translocate, such as those tested here, the agrochemical accumulates in the leaf Tips. By testing both leaf Tip and Middle, the relative amount of agrochemical moved by phloem or xylem transport since the leaf sprouted can be measured (Tip) along with the agrochemical in the process of transportation during harvest (Middle).

### 5.7 Pest Stage: Setup

Sections of maize leaves were placed in containers (INFO 3 x 6 x 2.5 cm deep) prepared with 5 mL of cooled 0.8% water agar to keep the container humidified along with a second instar *Spodoptera frugiperda*. Containers were sealed and placed in a dark chamber at 25 °C. After 4 d the containers were removed, and the contents evaluated. Leaf segments were graded based on feeding damage (e.g., 100% damage means no leaf remaining), while the FAW were evaluated for mortality (Figure 5C). Dead or moribund (dying) larvae were both recorded as dead.

### 5.8 Pest Stage: Analysis

Out of the anticipated 960 plants x 6 leaf segments per plant = 5760 assay wells, 49 individual wells were found to contain a FAW larvae that was dead upon infestation or the well contained no larvae. These observations are dropped resulting in 5711 total observations in the data set.

Feeding damage to each plant from the pest stage was compared across treatments according to three levels of granularity: overall feeding damage, feeding damage grouped by leaf (feeding damage separated into Leaf 3, Leaf 4, and Leaf 5), and by leaf segment (feeding damage separated into Leaf 3 Tip, Leaf 3 Middle, Leaf 4 Tip, Leaf 4 Middle, Leaf 5 Tip, and Leaf 5 Middle). Statistical significance was determined using pairwise t-tests with Bonferroni adjustment for the multiple hypothesis tests. To control for possible confounds of experimental run or tray number, feeding damage was regressed on treatment dummies as well as dummy variables for experimental run and tray location for overall feeding damage, FAW mortality rate, and feeding damage individually for each leaf segment (Tables SI 1 to SI 4). Tray number was significant due to selection method – tray 3 contained any less developed plants needed to fill the 48-plant total. All statistical analysis, including regressions and t-tests, were performed using Stata 17.0. Raw data and Stata code are available as supplemental materials.

## 6 Data Availability

Datasets related to this article can be found at https://osf.io/ykjsc/?view_only=790a53f9ed0d42639b3eef2677a91cd0, open-source online data repository hosted at Open Science Framework (28).

## 7 Supplemental Material

**Table SI 1:**
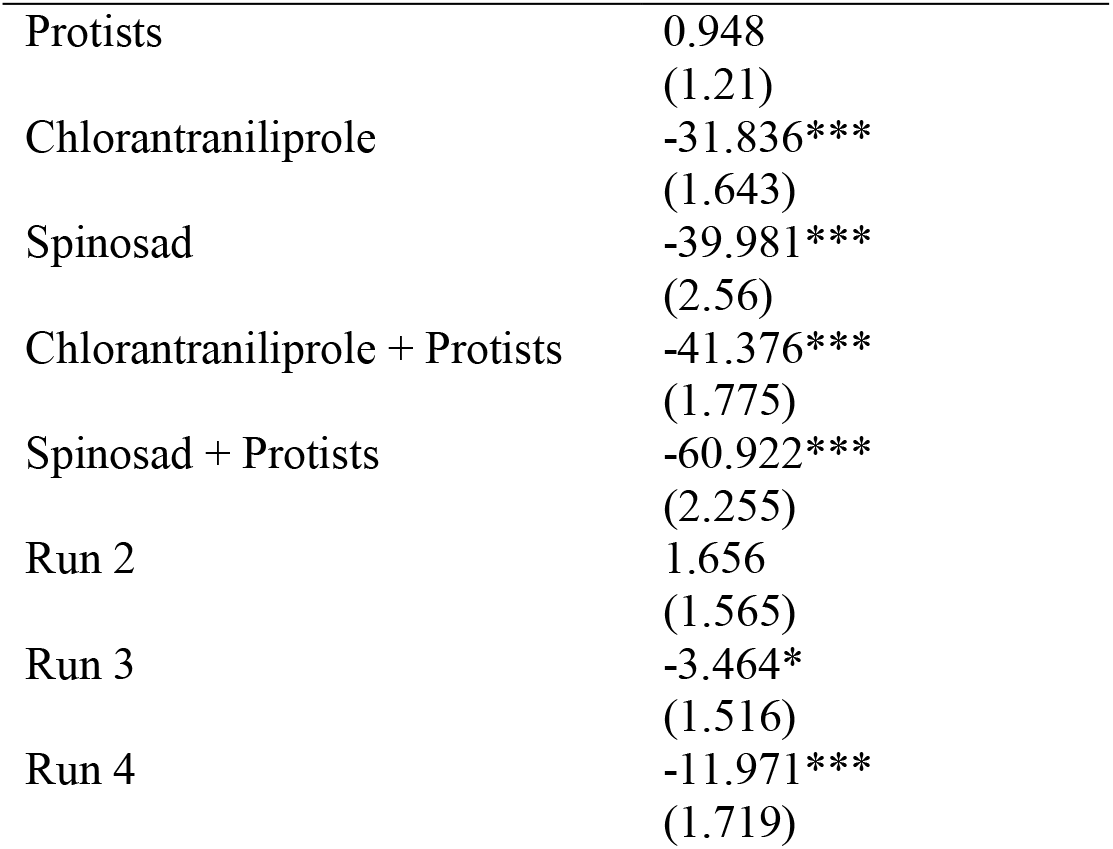
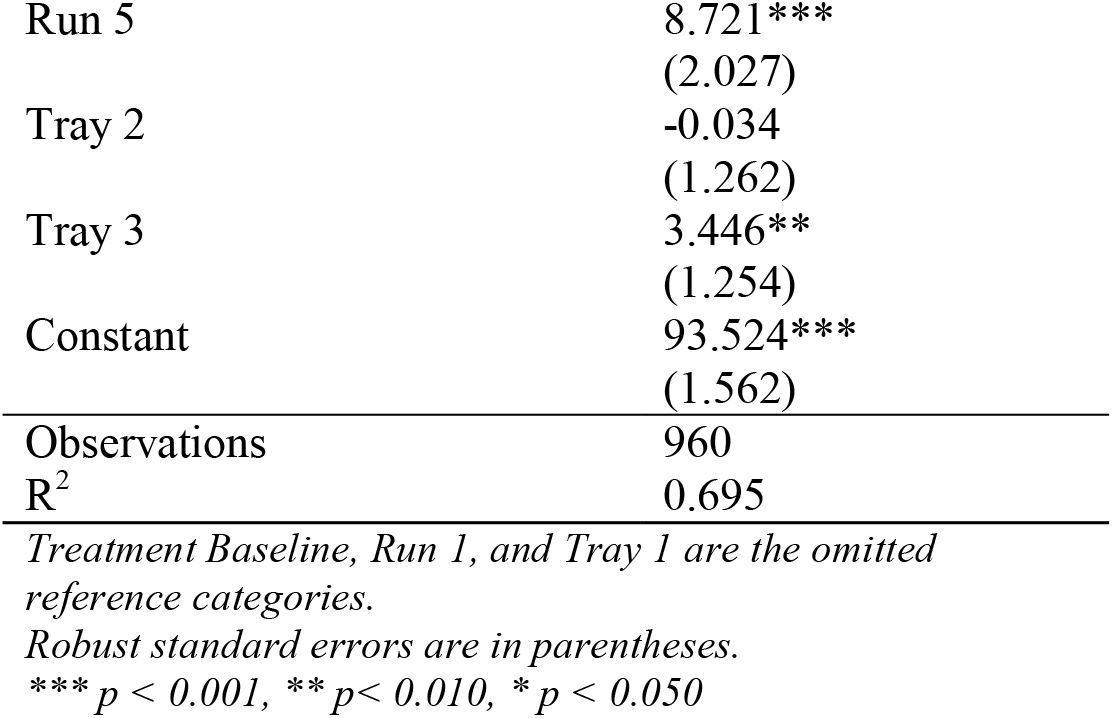
Regression for overall feeding damage (averaged across six leaf segment measures for each plant) with controls for experimental run and tray location

|  |  |
| --- | --- |
| Protists | 0.948<br>(1.21) |
| Chlorantraniliprole | -31.836***<br>(1.643) |
| Spinosad | -39.981***<br>(2.56) |
| Chlorantraniliprole + Protists | -41.376***<br>(1.775) |
| Spinosad + Protists | -60.922***<br>(2.255) |
| Run 2 | 1.656<br>(1.565) |
| Run 3 | -3.464*<br>(1.516) |
| Run 4 | -11.971***<br>(1.719) |
| Run 5 | 8.721***<br>(2.027) |
| Tray 2 | -0.034<br>(1.262) |
| Tray 3 | 3.446**<br>(1.254) |
| Constant | 93.524***<br>(1.562) |
| Observations | 960 |
| R<sup>2</sup> | 0.695 |

*Treatment Baseline, Run 1, and Tray 1 are the omitted reference categories.*
*Robust standard errors are in parentheses.*
\*\*\* $p < 0.001$ , \*\* $p < 0.010$ , \* $p < 0.050$

**Table SI 2:** Regression for FAW mortality rate (across six leaf segments for each plant) with controls for experimental run and tray location

|  |  |
| --- | --- |
| Protists | -0.937<br>(1.304) |
| Chlorantraniliprole | 20.341***<br>(1.685) |
| Spinosad | 33.342***<br>(2.642) |
| Chlorantraniliprole + Protists | 29.994***<br>(1.948) |
| Spinosad + Protists | 53.915***<br>(3.053) |
| Run 2 | -2.969<br>(1.665) |
| Run 3 | -2.552<br>(1.578) |
| Run 4 | 10.752***<br>(1.963) |
| Run 5 | -15.029***<br>(2.229) |
| Tray 2 | -0.042<br>(1.444) |
| Tray 3 | -2.833*<br>(1.398) |
| Constant | 5.605***<br>(1.634) |
| Observations | 960 |
| R<sup>2</sup> | 0.567 |

*Treatment Baseline, Run 1, and Tray 1 are the omitted reference categories.*
*Robust standard errors are in parentheses.*
\*\*\* $p < 0.001$ , \*\* $p < 0.01$ , \* $p < 0.050$

**Table SI 3:** Regression for feeding damage by leaf segment in Baseline, Protists, Chlorantraniliprole, and Chlorantraniliprole + Protists treatments with controls for experimental run and tray location

|  | (1)<br>Leaf 3<br>Middle | (2)<br>Leaf 3<br>Tip | (3)<br>Leaf 4<br>Middle | (4)<br>Leaf 4<br>Tip | (5)<br>Leaf 5<br>Middle | (6)<br>Leaf 5<br>Tip |
| --- | --- | --- | --- | --- | --- | --- |
| Protists | 0.015<br>(1.398) | 0.349<br>(1.398) | 0.401<br>(1.528) | 1.204<br>(1.811) | -0.760<br>(1.191) | 1.645<br>(1.056) |
| Chlorantraniliprole | -36.100***<br>(2.972) | -65.841***<br>(2.787) | -6.424**<br>(2.124) | -46.026***<br>(3.095) | -2.566<br>(1.500) | -34.783***<br>(2.978) |
| Chlorantraniliprole<br>+ Protists | -34.183***<br>(3.066) | -72.920***<br>(2.423) | -27.549***<br>(2.749) | -56.463***<br>(3.209) | -26.343***<br>(2.390) | -33.057***<br>(2.964) |
| Run 2 | -1.161<br>(2.857) | 9.866***<br>(2.349) | -6.856**<br>(2.347) | -13.021***<br>(3.117) | 10.990***<br>(2.096) | 11.382***<br>(2.556) |
| Run 3 | -4.312<br>(2.721) | 6.557**<br>(2.291) | -16.964***<br>(2.230) | -18.266***<br>(2.914) | 6.949***<br>(1.883) | 5.711*<br>(2.671) |
| Run 4 | -11.027***<br>(2.701) | 2.812<br>(2.448) | -15.203***<br>(2.300) | -10.762***<br>(2.806) | 3.068<br>(1.801) | 6.359***<br>(1.889) |
| Run 5 | 2.198<br>(2.158) | 8.781***<br>(1.860) | -5.201<br>(2.710) | -10.429**<br>(3.181) | 12.817***<br>(2.121) | 6.683**<br>(2.295) |
| Tray 2 | 0.087<br>(2.088) | 1.125<br>(1.649) | 1.984<br>(1.957) | -2.984<br>(2.246) | 4.756**<br>(1.493) | -0.003<br>(2.067) |
| Tray 3 | 3.980<br>(2.151) | 6.648***<br>(1.942) | 6.400***<br>(1.795) | 6.053**<br>(2.290) | 2.856<br>(1.556) | 3.101<br>(1.962) |
| Constant | 96.201***<br>(2.487) | 88.194***<br>(2.272) | 97.958***<br>(2.290) | 103.981***<br>(2.935) | 82.277***<br>(1.720) | 89.305***<br>(2.432) |
| Observations | 761 | 761 | 762 | 763 | 757 | 757 |
| R<sup>2</sup> | 0.347 | 0.734 | 0.271 | 0.519 | 0.321 | 0.381 |

*Treatment Baseline, Run 1, and Tray 1 are the omitted reference categories.*
*Robust standard errors are in parentheses.*
\*\*\* $p < 0.001$ , \*\* $p < 0.010$ , \* $p < 0.050$

**Table SI 4:** Regression for feeding damage by leaf segment in Baseline, Protists, Spinosad, and Spinosad + Protists treatments with controls for experimental run and tray location

|  | (1)<br>Leaf 3<br>Middle | (2)<br>Leaf 3<br>Tip | (3)<br>Leaf 4<br>Middle | (4)<br>Leaf 4<br>Tip | (5)<br>Leaf 5<br>Middle | (6)<br>Leaf 5<br>Tip |
| --- | --- | --- | --- | --- | --- | --- |
| Protists | -0.132<br>(1.497) | 0.188<br>(1.438) | 0.420<br>(1.487) | 1.228<br>(1.676) | -0.803<br>(1.245) | 1.890<br>(1.266) |
| Spinosad | -45.225***<br>(3.180) | -59.140***<br>(3.558) | -40.880***<br>(3.443) | -59.365***<br>(3.568) | -10.627***<br>(2.739) | -29.736***<br>(3.696) |
| Spinosad<br>+ Protists | -70.136***<br>(2.807) | -83.885***<br>(2.263) | -61.130***<br>(2.994) | -76.709***<br>(2.866) | -32.450***<br>(3.435) | -47.455***<br>(3.717) |
| Run 2 | 1.656<br>(2.859) | 3.635<br>(2.581) | -1.561<br>(2.572) | -0.198<br>(3.099) | 3.756<br>(2.100) | 5.361***<br>(1.594) |
| Run 3 | 4.187<br>(2.377) | 3.667<br>(2.570) | -4.323<br>(2.366) | 4.873<br>(2.587) | 2.421<br>(1.508) | 3.865<br>(2.001) |
| Run 4 | -17.522***<br>(2.887) | -8.321**<br>(2.718) | -11.335***<br>(2.555) | -3.336<br>(2.843) | -8.054***<br>(1.895) | -8.989***<br>(2.320) |
| Run 5 | 16.662***<br>(2.661) | 14.237***<br>(2.493) | 2.848<br>(2.762) | 6.065*<br>(3.051) | 16.057***<br>(2.054) | 16.246***<br>(2.449) |
| Tray 2 | -1.617<br>(1.798) | 0.012<br>(1.834) | -0.394<br>(1.984) | 0.056<br>(2.095) | 1.138<br>(1.833) | -1.812<br>(2.111) |
| Tray 3 | -0.453<br>(1.917) | -1.671<br>(1.885) | 0.677<br>(1.858) | 1.033<br>(1.957) | -1.162<br>(1.724) | 0.627<br>(1.869) |
| Constant | 94.548***<br>(2.463) | 94.459***<br>(2.511) | 94.685***<br>(2.430) | 92.669***<br>(2.831) | 88.790***<br>(1.543) | 93.210***<br>(2.211) |
| Observations | 664 | 666 | 666 | 669 | 666 | 664 |
| R<sup>2</sup> | 0.712 | 0.758 | 0.618 | 0.697 | 0.378 | 0.496 |

*Treatment Baseline, Run 1, and Tray 1 are the omitted reference categories.*
*Robust standard errors are in parentheses.*
\*\*\* $p < 0.001$ , \*\* $p < 0.010$ , \* $p < 0.050$

### 7.1 ^14^C Experiments

A scaled-up version of the µ-rhizoslide, described previously (18), was developed to work with maize. First, ^14^C-labeled agrochemical was dissolved in acetone, spiked onto the wall of a glass sample vial, and the acetone evaporated. Un-radiolabeled agrochemical formulation was then added to the vial and sonicated for 20 min to distribute the ^14^C-labeled material into the formulation. The formulation was then mixed with Page’s Saline containing heat-killed E. coli, or with Page’s Saline containing both protists and heat-killed E. coli to prepare the seed treatments formulation. As described before, the ^14^C-labeled seed treatment formulations were allowed to rest in darkness for 24 h prior to application. Final radioactivity was measured at approximately 300,000 DPM per individual seed treatment. Pots were filled, and seeds planted and treated as described above. Plants were harvested at 1 week due to restricted space for work with radiolabeled material. Once harvested, stem and leaves (stem segment) were separated from the roots and soil (root segment). Stem segments were freeze dried, then exposed on plates (20 x 25 mm Exposure cassette, Cytivia) for 24 h before being imaged by autoradiography (GE Amersham Molecular Dynamics Typhoon 9410 Molecular Imager). Plate images were processed using Image Quant TL software. The root segments had the seed remnants removed from the roots to prevent over-exposure on the plate due to being the location of initial dosing. Root segments then followed the same procedure as the stem portion of the plant. Plates for all treatments were imaged simultaneously to improve consistency within each experiment.

### 7.2 Transport of ^14^C-labeled Chlorantraniliprole

A similar experiment was conducted using ^14^C-labeled Chlorantraniliprole, differing in the allocated growth period (7 vs 14 d), watering schedule (due to limitations of radioactive containment), and pot (3D printed device optimized for radio-detection vs traditional 4.5” pot). Radioactivity in both the plant material and the soil were measured before combustion analysis was conducted on the plant material.

**Figure SI 1.**
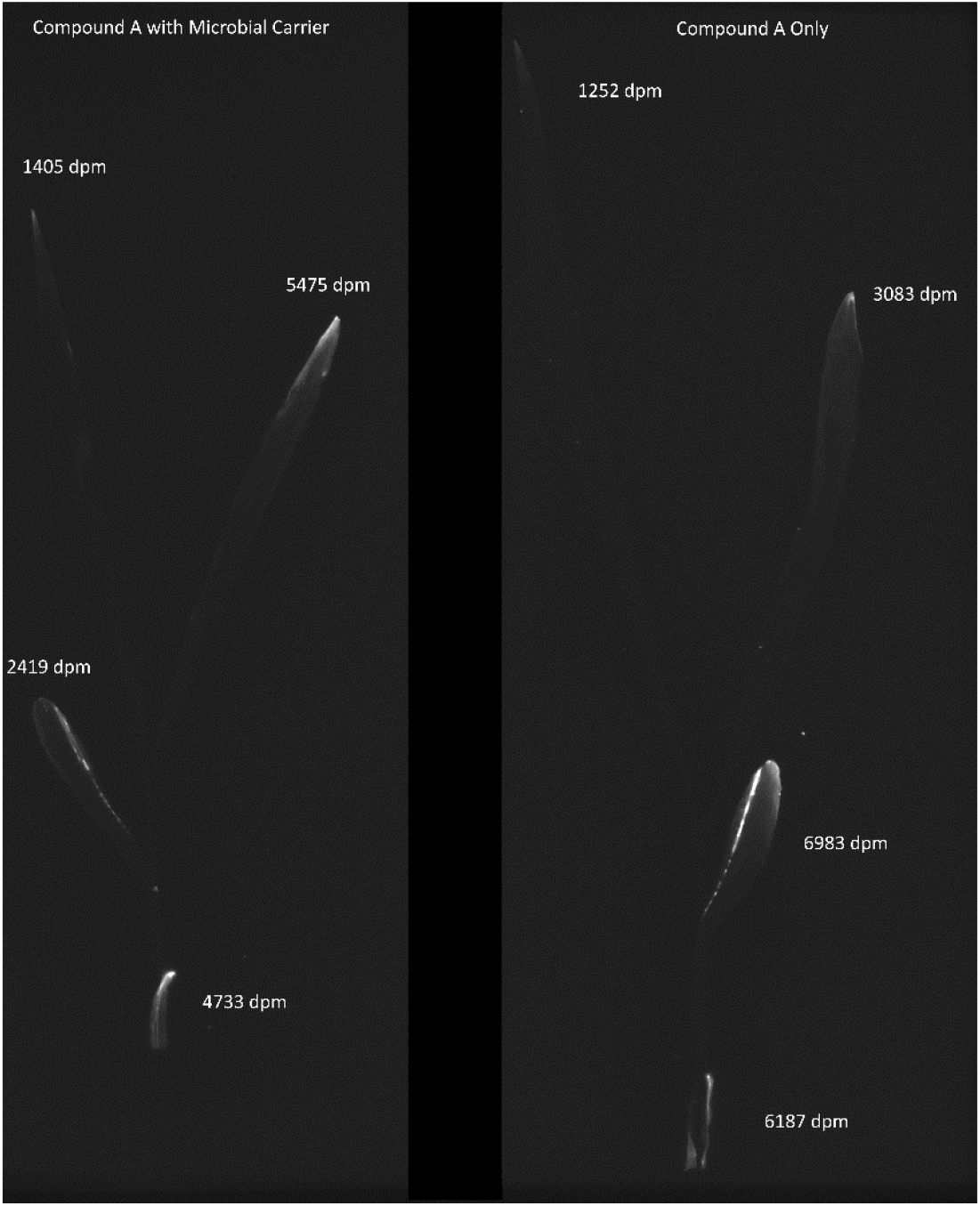
Radio image of plant matter from ^14^C experiment containing A) Chlorantraniliprole with protists, and B) Chlorantraniliprole alone. Each section is labelled with combustion analysis results –figure shows from the bottom up: stem, leaf 1, leaf 2, and leaf 3.

While the ^14^C results are too limited on their own to draw any definitive conclusions, they may show differing distribution of Chlorantraniliprole in the presence of protists when compared with the absence of protists, Figure 2. From these results, we see a general trend of higher concentration of Chlorantraniliprole in earlier parts (stem, Leaf 1) for Chlorantraniliprole alone when compared with Chlorantraniliprole and protists, and the reverse in later parts (Leaf 2, Leaf 3).

## 8 Acknowledgements

This work was supported by Corteva Agriscience, and NSF SusChEM award 1605816 (to LMS and DJG) and INTERN supplement, and USDA National Institute of Food and Agriculture award 2016-67013-24412 (to DJG and LMS).

